# Identification of two independent *COL5A1* variants in dogs with Ehlers Danlos syndrome

**DOI:** 10.1101/660407

**Authors:** Anina Bauer, John F. Bateman, Shireen R. Lamande, Eric G Hanssen, Shannon G.M. Kirejczyk, Mark Yee, Ali Ramiche, Vidyha Jagannathan, Monika Welle, Tosso Leeb, Fiona L Bateman

## Abstract

**BACKGROUND:** The Ehlers Danlos syndromes (EDS) are a heterogeneous group of heritable disorders affecting connective tissues. The mutations causing the various forms of EDS in humans are well characterized, but the genetic mutations causing EDS-like clinical pathology in dogs are not known, thus hampering accurate clinical diagnosis.

**RESULTS:** Clinical analysis of two independent cases of skin hyperextensibility and fragility, one with pronounced joint hypermobility was suggestive of EDS. Whole genome sequencing revealed *de novo* mutations of *COL5A1* in both cases, confirming the diagnosis of the classical form of EDS. The heterozygous *COL5A1* p.Gly1013ValfsTer260 mutation characterized in case 1 introduced a premature termination codon and would be expected to result in α1(V) mRNA nonsense-mediated mRNA decay and collagen V haploinsufficiency. While mRNA was not available from this dog, biochemical analysis of the dermis suggested reduced collagen V in the dermis and ultrastructural analysis demonstrated variability in collagen fibril diameter and the presence of collagen aggregates, termed ‘collagen cauliflowers’, consistent with *COL5A1* mutations underlying classical EDS. In the second case DNA sequencing demonstrated a p.Gly1571Arg missense variant in the *COL5A1* gene. While samples were not available for further analysis, such a glycine substitution would be expected to destabilize the strict molecular structure of the collagen V triple helix and thus affect protein stability and/or integration of the mutant collagen into the collagen V/collagen I heterotypic dermal fibrils.

**CONCLUSIONS:** This is the first report of genetic variants in the *COL5A1* gene causing the clinical presentation of EDS in dogs. These data provide further evidence of the important role of collagen V in dermal collagen fibrillogenesis. Importantly from the clinical perspective we show the utility of DNA sequencing, combined with the established clinical criteria, in the accurate diagnosis of EDS in dogs.

## BACKGROUND

The Ehlers Danlos syndromes (EDS) are a heterogenous group of heritable disorders affecting connective tissue. In humans, variable clinical manifestations affecting primarily the skin, joints, ligaments, blood vessels and internal organs have been reported. EDS has also been seen in many animal species, including horses [1], mink [2], rabbits [3], dogs [4-14], and cats [15-17]. Whilst a rare condition in dogs, it has been most frequently reported in Dachshunds, Boxers, St. Bernard’s, Greyhounds, Irish Setters and Poodles [18].

Abnormal collagen fibril formation is the hallmark of several types of human EDSs including the classical form [19]. In the classical form of EDS, mutations in *COL5A1* and *COL5A2*, which encode the α1- and α2-chain of type V collagen, have been reported [20-26]. Type V collagen is a quantitatively minor fibril-forming collagen. Several isoforms exist, but the most widely distributed form is the [α1(V)]_2_ α2(V)] heterotrimer that co-assembles with type I collagen into heterotypic type I/V collagen fibrils in the extracellular matrix. Type V collagen is thought to regulate the diameter of these fibrils by retention of its large N-propeptide domain, which projects above the surface of the collagen fibril [27]. In humans, mutations in *COL5A1* and *COL5A2* lead to characteristic collagen abnormalities in the skin with variability in collagen fibril diameter and the presence of collagen aggregates, termed ‘collagen cauliflowers’ [28]. Abnormal collagen structure and fibril formation in turn contribute to the clinical signs seen in classic EDS including hyperextensible skin, generalized joint hypermobility and generalized connective tissue fragility [29].

We report here several dogs with genetic variants in the *COL5A1* gene affecting collagen synthesis and structure, analogous to classical EDS in human patients. Furthermore, we describe histopathological and ultrastructural findings associated with the *COL5A1* variants in these animals. Collagen protein analysis of one affected individual was also performed.

### Methods

#### Ethics

Blood and skin samples collected from case 1 and the related controls (FB and MY) were approved by the Murdoch Children’s Research Institute Animal Ethics Committee (Approval # A815). Case 1 was humanely euthanized at the request of the owner immediately prior to sample collection. This was performed using IV pentobarbitone as per standard techniques. Blood and skin samples for cases 2 and 3 (RA) were collected as part of routine veterinary diagnostic assessment. Taking of blood samples from healthy control dogs for genetic analysis was approved by the “Cantonal Committee For Animal Experiments” (Canton of Bern; permit 75/16).

### Histopathology

#### Case 1

One 6 mm diameter punch biopsy was obtained from the dorsum of the 6-month-old Labrador, fixed in 10 % neutral-buffered formalin, processed routinely and embedded in paraffin wax. Four-to-five-micron-thick sections were stained with hematoxylin and eosin (H&E), Masson’s Trichrome (MT), and Verhoeff van Gieson (VVG) stains at the University of Georgia, College of Veterinary Medicine Histology Laboratory. As a control, a punch biopsy was obtained from the abdominal region of a 10-month-old, intact male, Labrador dog, fixed in formalin, processed routinely, and stained with H&E, MT, and VVG for comparison. Slides were reviewed by a board certified veterinary pathologist (SK).

#### Cases 2 and 3

One 8 mm punch biopsy from the lateral thorax was investigated histologically. The biopsy was fixed in 10 % buffered formalin and processed routinely. The slide was stained with hematoxylin and eosin (H&E) prior to histopathological examination by a board certified veterinary pathologists (MW).

##### Transmission Electron Microscopy (TEM)

For electron microscopy dermal samples were fixed in 0.1 M sodium cacodylatecontaining 2.5 % glutaraldehyde, after rinsing the samples were post-fixed in 1 % aqueousosmium tetroxide, dehydrated in an alcohol series, and embedded in Epon 812. Seventy nanometer ultrathin sections were cut and observed on a Tecnai F30 with an extraction voltage of 200 kV. Micrographs were taken using a Gatan UltraScan 1000.

##### Biochemical analysis of dermis

Full thickness dermal biopsies were diced with a scalpel and defatted by gentle shaking with cold chloroform:methanol (2:1) for 24 hrs. Tissue was washed with cold methanol and dried under vacuum. Dried tissue was re-hydrated in 50 mM Tris/HCl, pH 7.5 containing 0.15M NaCl. To assist in the uniformity of extraction, the re-hydrated tissue was snap frozen and powered under liquid nitrogen. For collagen analysis a sequential extraction protocol was used to extract the successively more cross-linked collagen matrix as previously described [30]. The freeze-milled dermis was first extracted with 0.15 M NaCl, 50 mM Tris-HCl buffer for 24 h at 4°C to remove the soluble collagens followed by extraction with 4 M GuHCl (Tris-HCl buffer [pH 7.4]), then 0.5M acetic acid, and finally digestion with pepsin at 100 μg/ml in 0.5M acetic acid to sequentially extract the successively more cross-linked and thus mature collagen matrix. To enrich for collagen V, selective salt precipitation of portions of the pepsin extracted collagen was performed [31]. Collagen chains in each extract were analyzed on SDS–gradient polyacrylamide gels (3-8%, Novex), visualized by Coomassie Brilliant Blue staining and quantified as described previously [30]. Pepsin-soluble collagen from rat tail tendon was run on each gel as a standard for quantitation.

### DNA extraction and whole genome sequencing

Genomic DNA was isolated from EDTA blood samples of the investigated dogs. For case 1 and case 2, an Illumina PCR-free TruSeq fragment library with ∼390 bp insert size was prepared. We collected ∼153 and 211 million 2 × 150 bp paired-end reads or ∼18 × and ∼25 × coverage on an illumina HiSeq 3000 instrument for cases 1 and 2, respectively. The reads were mapped to the CanFam3.1 dog reference genome assembly and aligned using Burrows-Wheeler Aligner (BWA) version 0.7.5a [32] with default settings. The generated SAM file was converted to a BAM file and the reads were sorted by coordinate using samtools [33]. Picard tools (http://sourceforge.net/projects/picard/) was used to mark PCR duplicates. To perform local realignments and to produce a cleaned BAM file, we used the Genome Analysis Tool Kit (GATK version 2.4.9, 50) [34]. GATK was also used for base quality recalibration with canine dbsnp version 139 data as training set. The sequence data were deposited under the study accession PRJEB16012 and sample accessions SAMEA104091568 (case 1), and SAMEA4867923 (case 2), at the European Nucleotide Archive. Additionally, we used 356 additional whole genome sequences as controls, which were either publicly available [35], produced during other projects of our group or contributed by members of the Dog Biomedical Variant Database Consortium.

Putative SNVs were identified in each of the 358 samples individually using GATK HaplotypeCaller in gVCF mode [36]. Subsequently all sample gVCF files were joined using Broad GenotypeGVCFs walker (-stand_emit_conf 20.0; -stand_call_conf 30.0). Filtering was performed using the variant filtration module of GATK using the following standard filters: SNPs: Quality by Depth: QD < 2.0; Mapping quality: MQ < 40.0; Strand filter: FS > 60.0; MappingQualityRankSum: MQRankSum < - 12.5; ReadPosRankSum < −8.0. INDELs: Quality by Depth: QD < 2.0; Strand filter: FS > 200.0. The functional effects of the called variants were predicted using SnpEFF software [37] together with the NCBI annotation release 105 on CanFam 3.1. Identified sequence variants were confirmed by Sanger sequencing of PCR amplicons. Targeted genotyping of additional dogs was also done by Sanger sequencing.

## RESULTS

### Clinical assessment - Case 1

A 4-month-old male intact Labrador was examined for several small skin lacerations of the distal limbs, hock and tail base. On physical examination the dog was found to be clinically healthy, with normal skeletal development. Several seroma-like swellings were noted in the regions of skin trauma (hocks, tail base). Aspirates taken from these swellings revealed moderate numbers of erythrocytes with fewer neutrophils and macrophages with proteinaceous debris. These findings were consistent with a seroma or hematoma formation and the dog was treated with amoxycillin/clavulanic acid (Clavulox®, Zoetis, Ryde, Australia) and firocoxib (Previcox®, Zoetis, Ryde, Australia).

Two months after initial presentation the dog was again examined for ongoing seroma formation and joint abnormalities. On physical examination evidence of generalized joint hyperextensibility (Figure 1) and skin hyperextensibility (Figures 2,3) and fragility was noted. The dog had a skin extensibility test [38] of 23.5 % (normal < 14.5; %).[39] A complete blood count revealed a mild anemia of 4.87×10^12^/L (5.5-8.5), HCT 0.33 L/L (0.37-0.55), Hb 113 g/L (120-180) and a normal white cell count. Serum biochemistry showed a mild hypoalbuminemia 21 g/L (28-42) and hypoproteinemia 47 g/L (54-78). Other serum abnormalities included a moderate hyperphosphatemia, mild elevation in ALP and decrease in creatinine which were all considered within normal limits for a young dog. At this time, due to the ongoing joint hypermobility and skin lacerations with the suspicion of a genetic collagen disease, the owners elected to humanely euthanize the dog.

**Figure 1.**
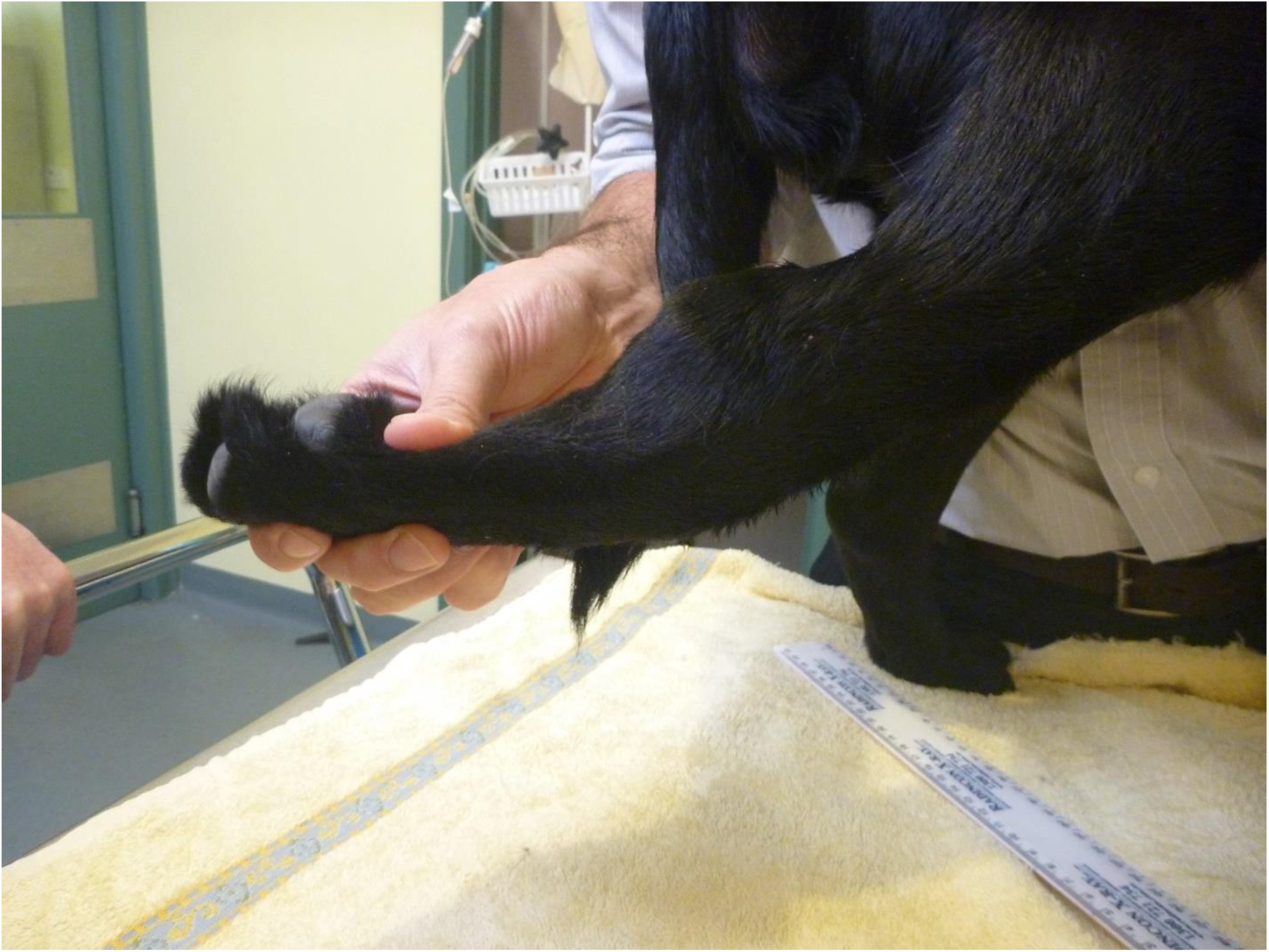
Case 1: Joint hypermobility of the tarsus and metatarsus.

**Figure 2.**
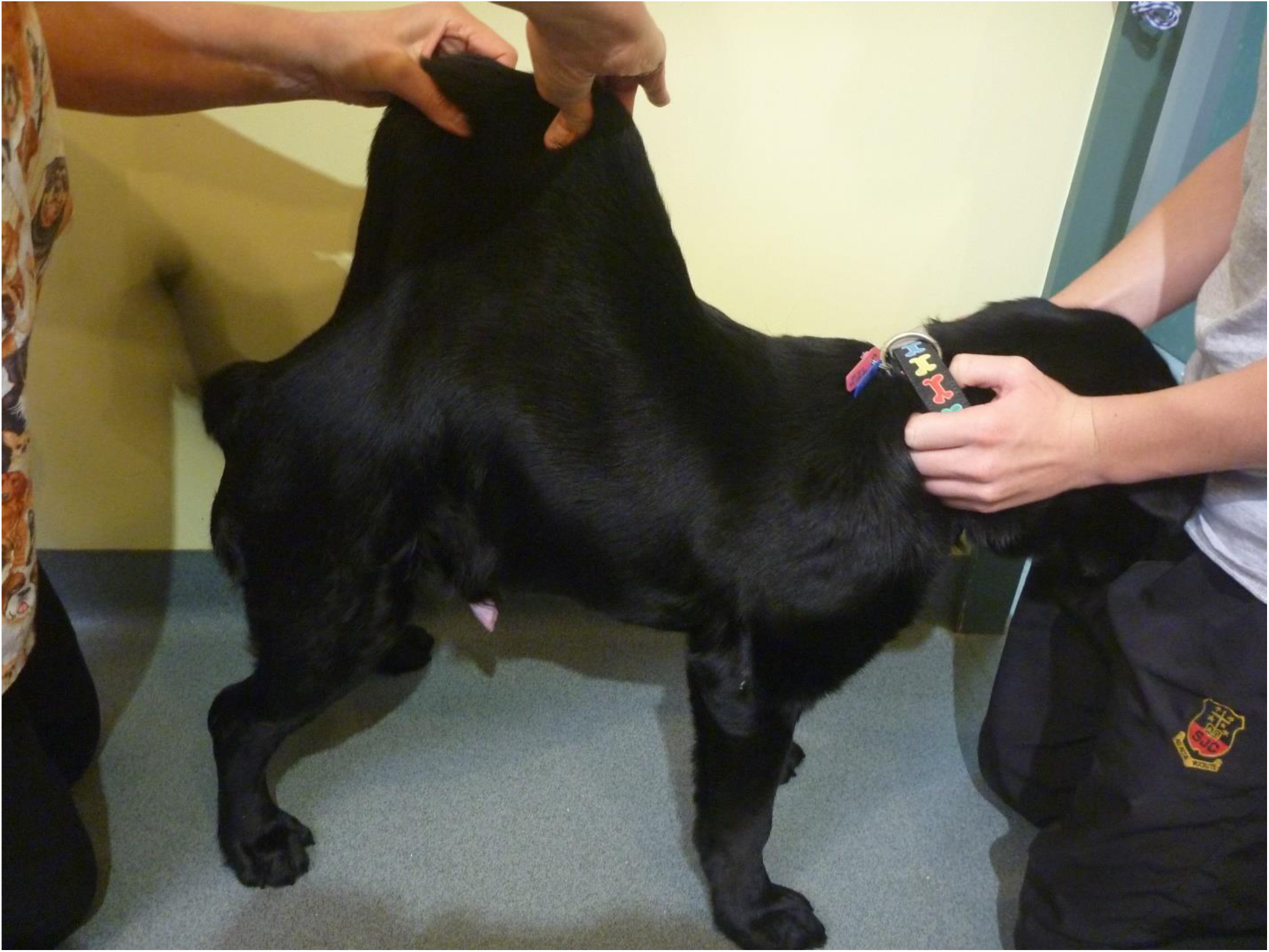
Case 1: Skin hyperextensibility over the dorsum.

**Figure 3.**
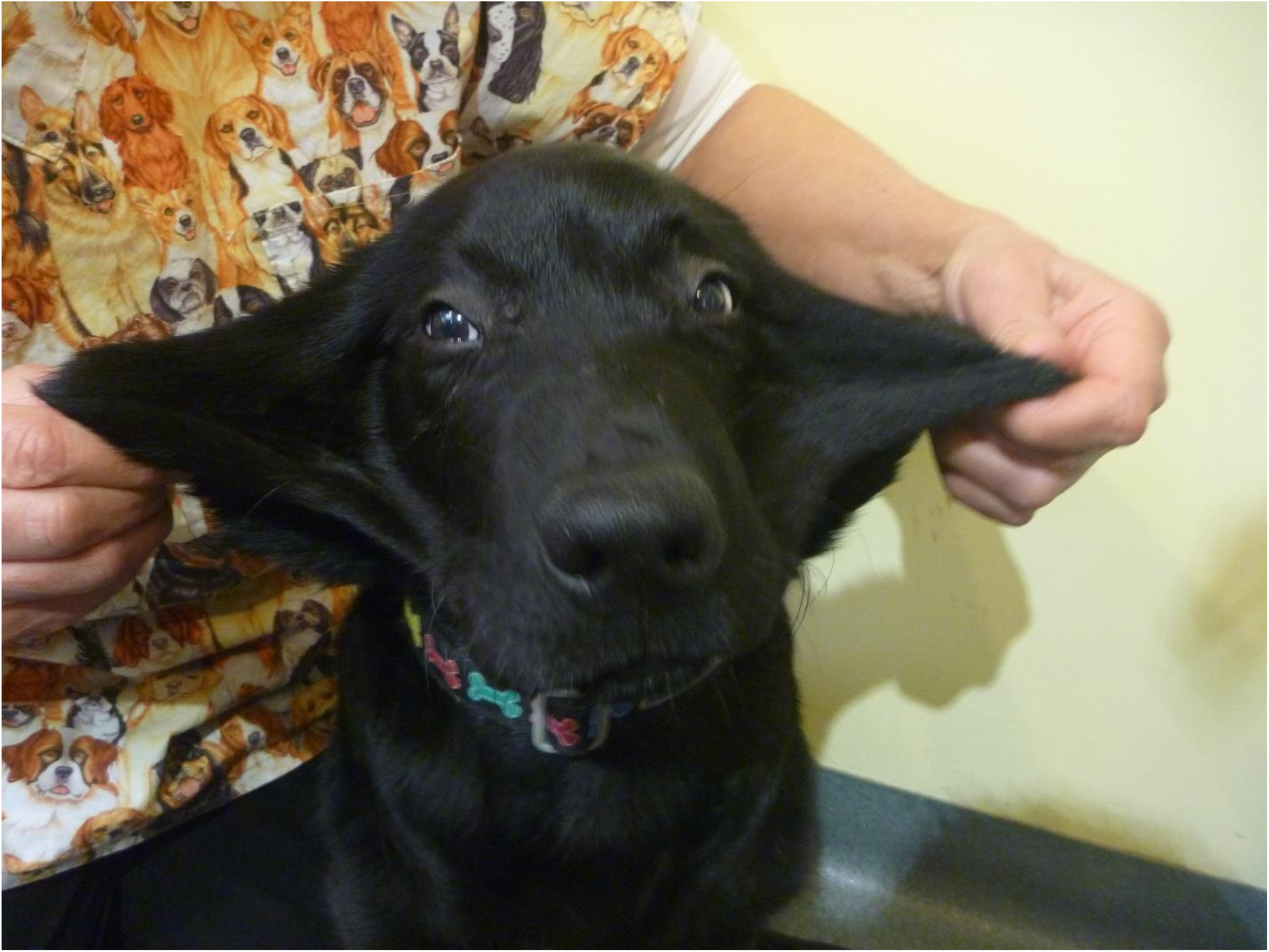
Case 1: Skin hyperextensibility of the face.

### Cases 2 & 3

Two young adult mixed breed dogs were presented for hematomas and skin lacerations after mild trauma. The dogs (siblings) were adopted from an animal shelter is the south east region of Morocco – the prior clinical history and inset of clinical signs were unknown, though it was noted that the dam was unaffected. On physical examination the dogs were found to be clinically healthy, with normal skeletal development. Bruising and skin lacerations were noted on the carpi with oozing, hemorrhagic zones of 3-4cm in size with disruption of the skin barrier (splitting of the skin) in these regions (case 2). Bruising and wounds were noted to occur after only mild trauma. There was hyperelasticity of the elbows (Figure 4) and tarsi as well as the skin of the neck (Figure 5) and flank (case 2 and 3), with evidence of previous scarring on the neck (case 2). The rest of the clinical exam was unremarkable for both dogs.

**Figure 4.**
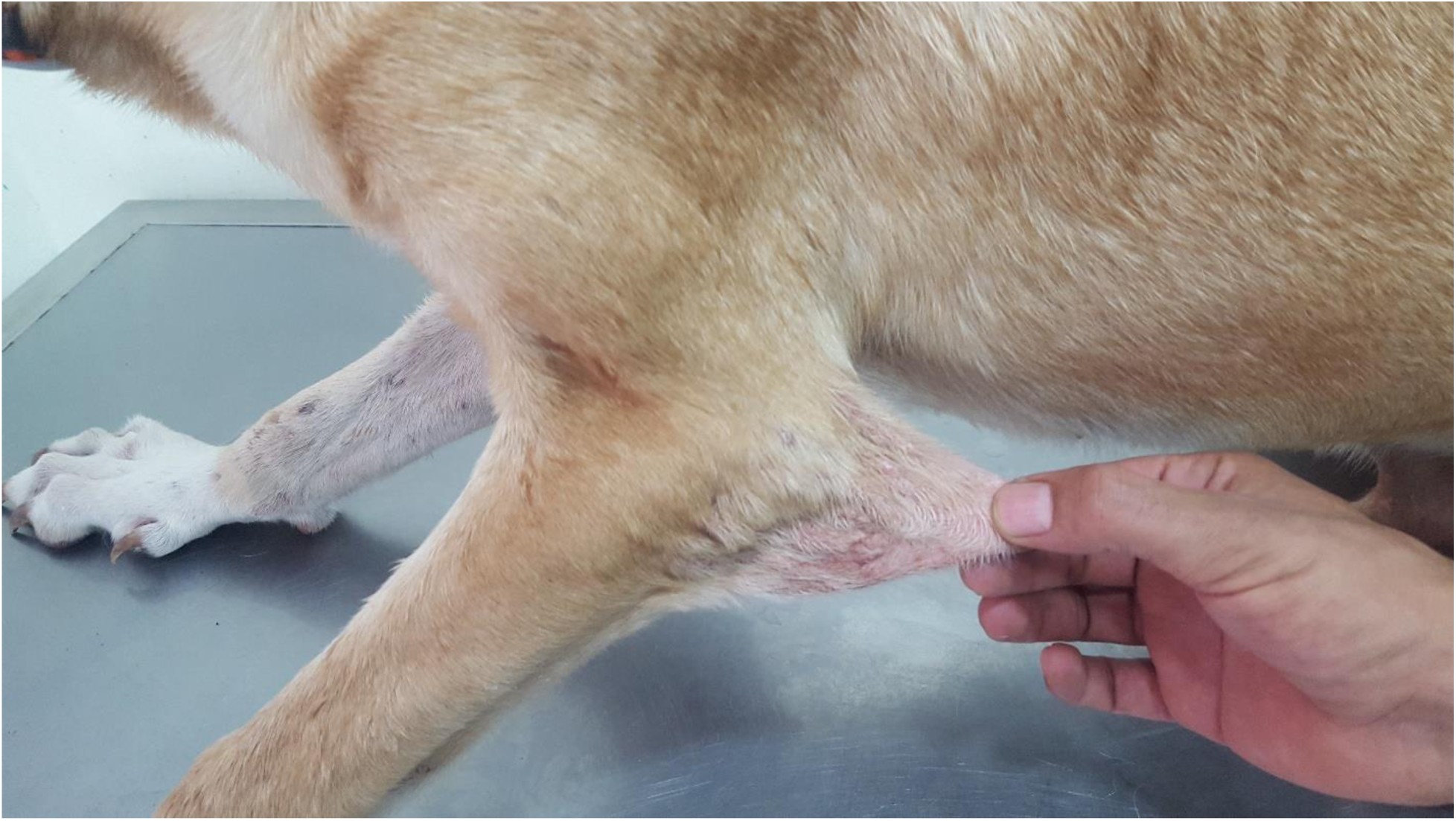
Case 2: Skin hyperextensibility over the elbow.

**Figure 5.**
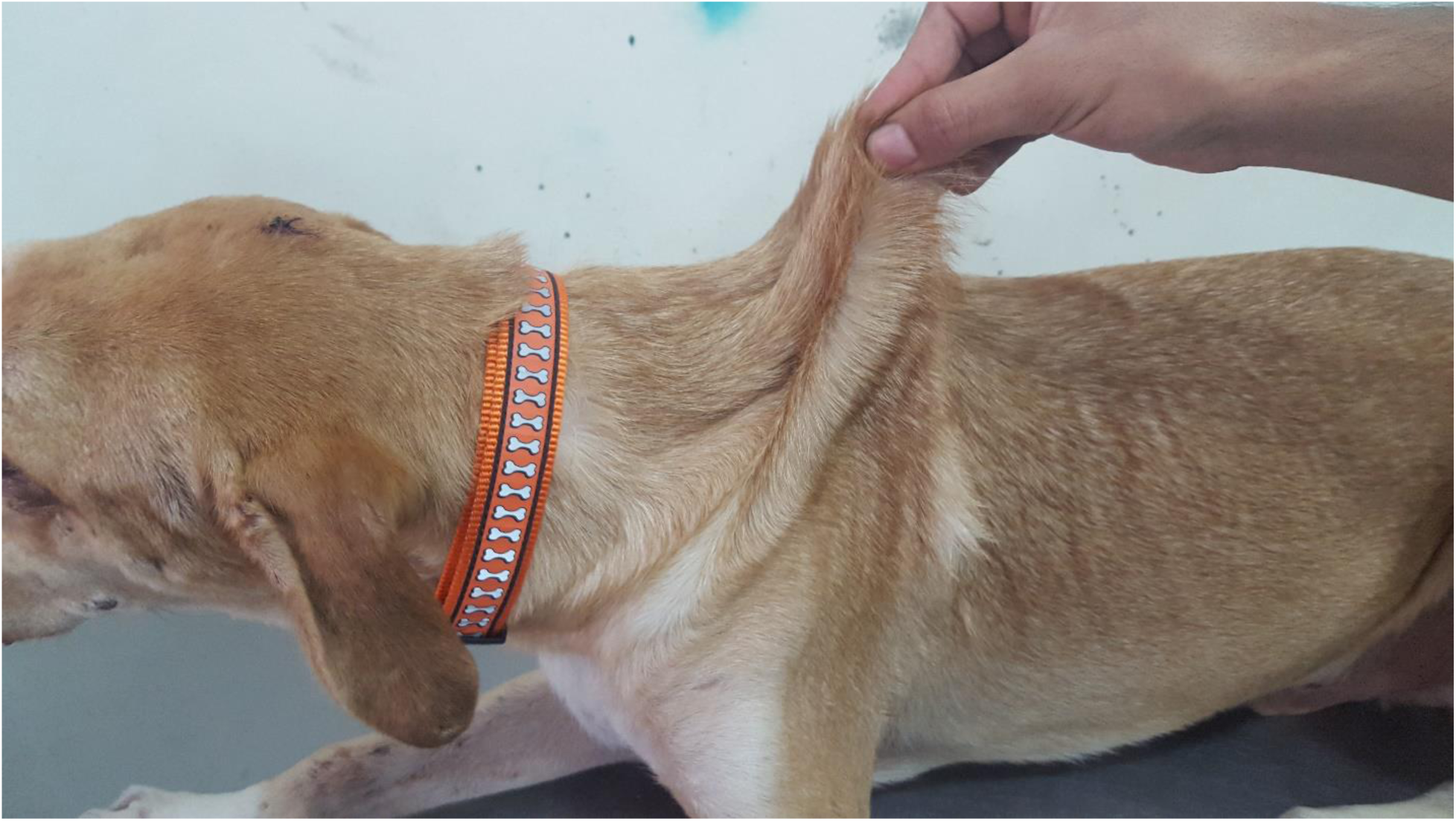
Case 2: Skin hyperextensibility over the dorsum.

On routine screening blood work for case 2, the hematocrit was normal at 0.48 L/L (0.37-0.55) and total solids 54 g/L (54-78). Whilst skin biopsy of case 2 and 3 were performed, no further diagnostic testing was undertaken. The outcome of these dogs is currently not known.

### Histopathology

#### Case 1

Histological examination of skin from the 4-month-old Labrador revealed dermal collagen fiber abnormalities, characterized by fiber disarray, variability in fiber diameter, and multifocal fiber clumping (Figure 6, Panels A & C). Collagen clumps stained hypereosinophilic on H&E and dark red on a Masson’s Trichrome stain. On the Verhoeff van Gieson stain, dermal elastic fibers were mildly increased in number multifocally, irregularly distributed throughout the dermis, and variably sized. The space between collagen fibers as well as the number and arrangement of dermal fibroblasts was similar between the affected and control dogs. The epidermis was of normal thickness in both the affected and control dog skin.

**Figure 6.**
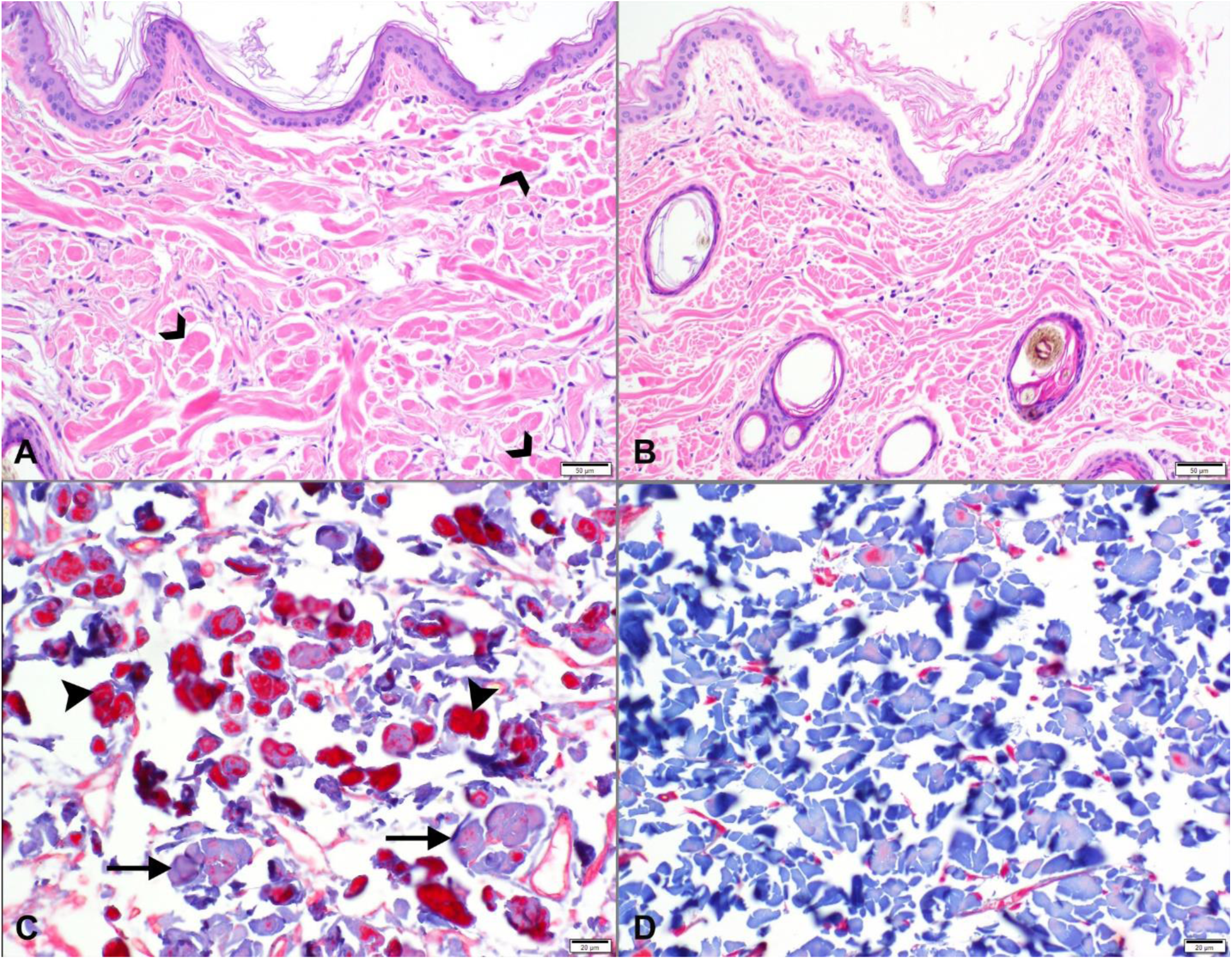
Histopathology of skin from a 4-month-old, male intact, Labrador retriever dog (Case 1) with EDS-like syndrome (Panels A and C) compared to skin from an unaffected, 10-month old, male intact Labrador retriever dog control (Panels B and D). A: Photomicrograph of skin from the dorsum of affected dog demonstrates dermal collagen fiber disorganization with variation in fiber diameter, length and staining intensity. Collagen fibers occasionally form variably sized, hypereosinophilic clumps (open arrowheads). 200× original magnification; bar = 50 microns. Hematoxylin and Eosin stain. B: The control dog skin has a more organized, often parallel dermal collagen fiber arrangement and fibers are of more uniform diameter. 200x original magnification; bar = 50 microns. Hematoxylin and Eosin stain. C: On the Masson’s Trichrome (MT) stain, clumped collagen fibers are variably sized, irregularly distributed throughout the dermis and often stain deep red (arrowheads). There are fewer, more normal collagen fibers, which stain pale blue on the MT stain (arrows). 400x original magnification; bar = 20 microns. D: Compared to panel C, the unaffected control dog’s superficial dermis exhibits mostly uniform, pale blue staining of collagen fibers on the MT stain. 400x original magnification; bar = 20 microns.

#### Cases 2 and 3

Histological examination of the skin biopsies from these affected siblings revealed abnormal collagen fibers characterized by a severe variability in fiber diameter ranging from thin and wispy to normal thickness. In addition there was shortening of the fibers and fiber disarray. Abnormal fibers were outlined by increased numbers of fibroblasts. Space between fibers was wider than normal and there was abundant hemorrhage especially in the mid dermis. The dermis and the overlying epidermis were of normal thickness.

### Transmission Electron Microscopy

The clinical and histological features discussed above suggested that all three cases may have a form of Ehlers Danlos syndrome. In case 1, tissue was available for detailed ultrastructural analysis and transmission electron microscopy of the dermis revealed abnormal fibril architecture which was also consistent with EDS. While in the control dermis the collagen fibrils were highly organized and aligned and with an approximately circular cross-sectional shape of uniform diameter (Figure 7, A-D), the affected dog dermis had in addition to this normal fibril architecture a significant second population of disorganized collagen fibrils with large and irregular cross-sectional shapes (Figure 7, E-H). This fibrillar disorganization has been widely reported in human EDS patient skin[20] and in mouse models of EDS[19] and has also been reported in dogs[11]. Unfortunately, samples were not available from cases 2 and 3 for ultrastructural analysis of collagen fibrillar architecture.

**Figure 7.**
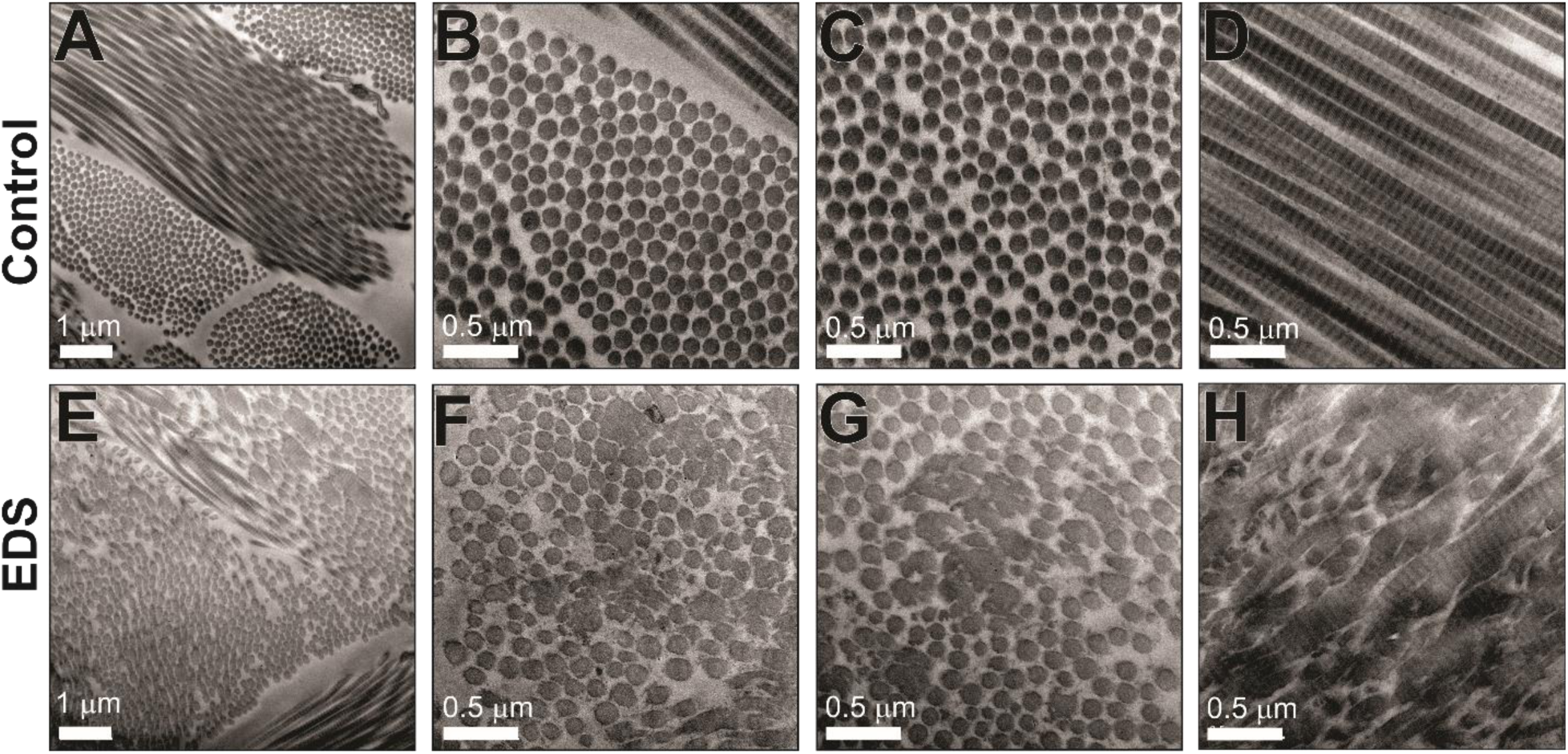
Structurally abnormal dermal collagen fibrils in Case 1. Transmission electron micrographs from the dermis of control (A-D) and Case 1 (EDS) (E-H). Control dermis contains well-organized collagen fibrils of uniform size and circular profile, while the EDS dermis contains two populations of fibrils. One fibril population is similar to the control dermis whereas the other population of fibrils are large and structurally abnormal. Scale bars of 0.5µm or 1µm are shown.

### Biochemical analysis of dermis

For case 1 dermal tissue was also available for biochemical analysis of the collagen composition. Extraction with isotonic neutral pH buffer (NSS), which removes the newly synthesized collagen species, accounted for only a small proportion of the total collagen extracted (Figure 8A, lanes 1-2) as did further extraction with GuHCl (lanes 3-4) or acetic acid (lanes 5-6) which extracts newly crosslinked collagen matrix. Note that lanes 1-6 were loaded with much higher proportions of the extracts to achieve Coomassie-detectable bands (Figure 8A, see legend for loading details). In all these extracts, collagen I and collagen III were the most abundant collagens with no apparent quantitative differences between EDS and control. The pepsin extracts of the dermis (Figure 8A, lanes 7-8) released the more crosslinked collagenous matrix as evidenced by the abundance of type I crosslinked dimers (β-components) and trimers (γ-components). There were no detectable qualitative difference in collagen I and III ratios or in collagen I crosslinking in these samples. However, while below the levels required for reliable quantitative analysis, the ratio of α1(V):α1(I) appeared to be reduced, suggestive of reduced collagen V in the EDS dermal samples. Collagen V was partially purified to provide additional qualitative data on the pepsin-extracted α1(V) and α2(V) chains (Figure 8B). Both α1(V) and α2(V) chains had a normal electrophoretic migration excluding major structural mutations affecting the α-chain size, or increased post-translational modifications which could indicate helix-disrupting mutations.

**Figure 8.**
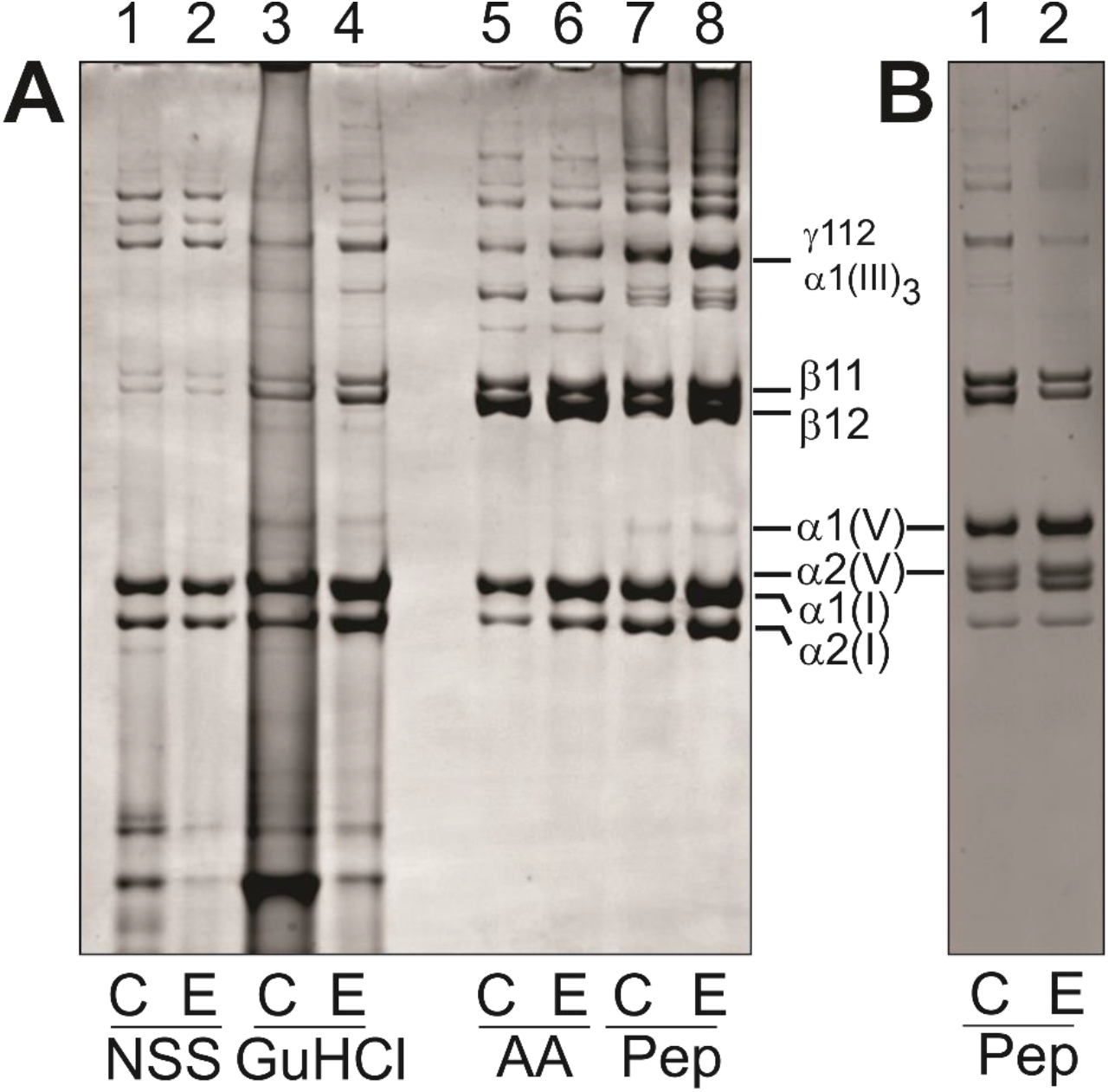
Dermal collagen extracellular matrix composition. Samples from control dermis (C) and EDS dermis (E) were extracted sequentially 0.15M NaCl, 50mM Tris-HCl buffer, pH7.4 (neutral salt soluble, NSS), 4M GuHCl (GuHCl), 0.5M acetic acid (AA) and digestion with pepsin (Pep) at 100 μg/ml to sequentially extract the successively more cross-linked and thus mature collagen matrix. **(A)** Extracts analyzed by SDS-polyacrylamide gel (3-8%) electrophoresis; samples unreduced. **(B)** Pepsin extracts where collagen V chains were enriched by selection salt precipitation. Note that all samples were loaded to achieve a comparable staining intensity of control and EDS sample pair in each extract and do not reflect the relative amount of collagen between extract. The vast majority of the collagen was found in the pepsin extract fractions (>95%) of both the Control and EDS dermis. The migrations of collagen V α1(V) and α2(V) and the collagen I α1(I) and α2(I) chains and crosslinked -chain dimers (β11 and β12) and trimers (γ112) are shown. The disulfide-bonded collagen III trimer is denoted by α1(III)_3_.

### Genetic analyses

#### Case 1

We sequenced the whole genome of the affected Labrador Retriever at ∼18x coverage and called SNVs and short indels with respect to the canine reference genome (CanFam 3.1). The variants were compared to 356 control dogs from diverse breeds. We analyzed the data under two alternative models regarding the mode of inheritance: monogenic recessive or monogenic dominant (and due to a *de novo* mutation event). For the recessive model we only considered variants, at which the affected dog was homozygous for the non-reference allele and none of the control dogs carried the non-reference allele. Hard-filtering resulted in 4 private protein-changing variants present in the affected dog (Table 1). None of these variants was located in a known candidate gene for Ehlers-Danlos syndrome. When filtering for heterozygous genotypes in the affected dog under the alternative dominant model, our automated pipeline detected 31 private protein-changing variants, among them a single nucleotide deletion in the *COL5A1* gene. This variant, XM_022423936.1:c.3038delG, leads to a shift in the reading frame and a premature stop codon p.(Gly1013ValfsTer260). The heterozygous genotype at this variant in case 1 were confirmed by Sanger sequencing, and the genotypes of the parents were determined (Figure 9). In the affected dog, the ratio of wildtype:mutant allele was ∼1:1. The sire of case 1 was homozygous wildtype. In contrast, the wildtype:mutant allele ratio was ∼4:1 in DNA isolated from blood of the affected dog’s dam, indicating that the mutation event had occurred in the dam and that she was a genetic mosaic.

**Figure 9.**
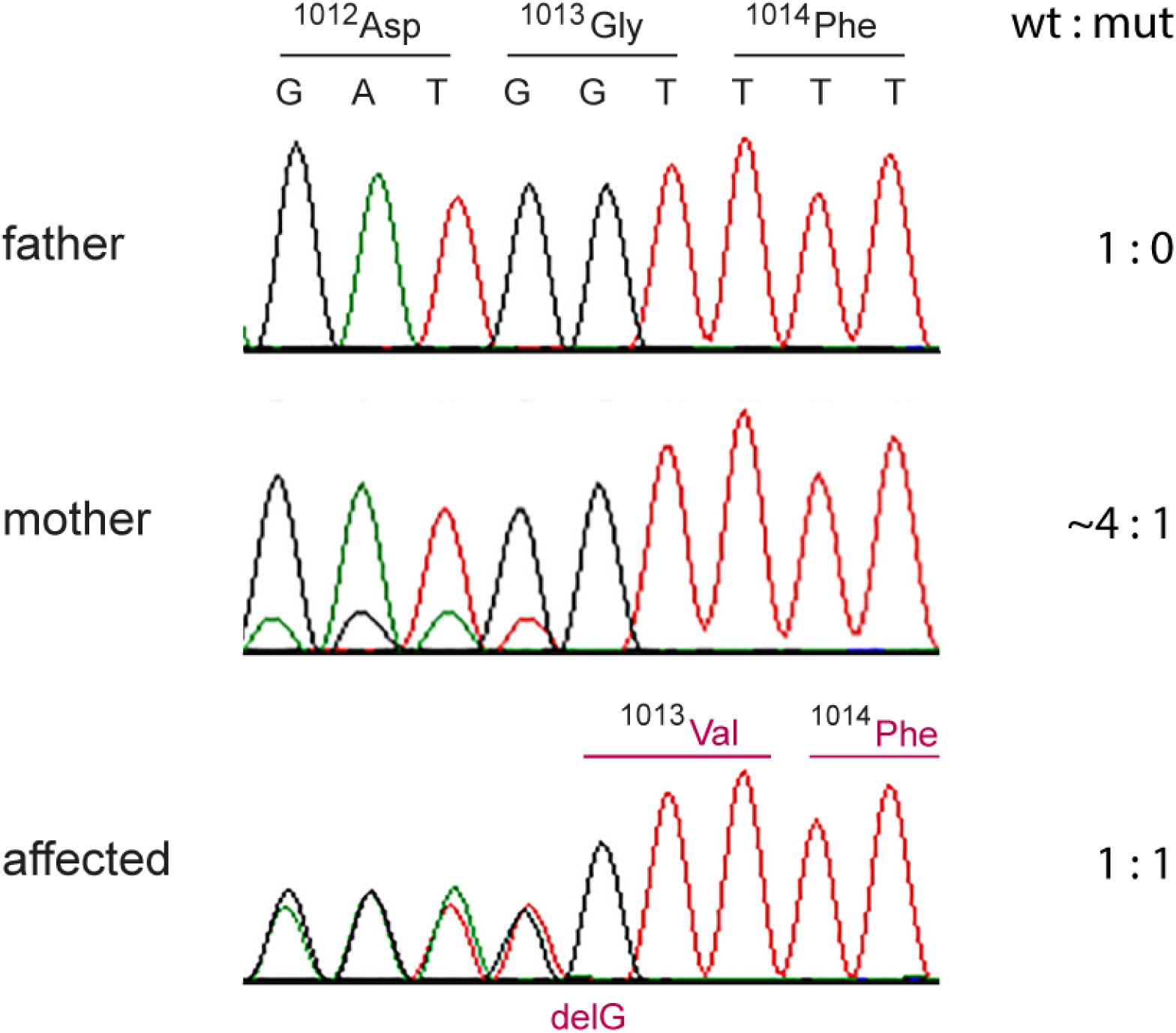
Sanger electropherograms illustrating the c.3038delG variant in case 1 and its parents. The sequence reads were obtained with a primer in reverse complementary orientation. While the affected dog shows a 1:1 ratio of the two alleles indicating a heterozygous genotype, the skewed allele ratio in the mother suggests genetic mosaicism. The mutant allele in the mother must have been present in white blood cells and in her germline. This confirms that the c.3038delG variant arose by a *de novo* mutation event during development of the mother.

#### Cases 2 and 3

We sequenced the whole genome of case 2 at ∼25× coverage. As in case 1, SNVs and short indel variants were called and compared to 356 control dogs, considering both a recessive and a dominant mode of inheritance. Our automated pipeline detected 9 homozygous protein-changing variants in the affected dog but none of them in a functional candidate gene (Table 1). Hard filtering for private protein-changing variants present in the heterozygous state in case 2 resulted in 276 variants, including a missense variant in the *COL5A1* gene, XM_022423936.1:c.4711G>A; XP_022279644.1:p.(Gly1571Arg). Sanger sequencing confirmed a heterozygous genotype at the variant in case 2 and the available affected full-sibling (case 3), and the absence of the variant in 493 dogs from 66 different breeds (Table S1). DNA from the parents was not available.

## DISCUSSION

The clinical presentation in all three cases was consistent with a form of EDS. The most common form of EDS in humans, classical EDS (cEDS) is an autosomal dominant condition [40]. In 2017, Malfait et al. proposed the International Classification for EDS [29]. With recent developments in next generation sequencing and the discovery of a number of genetic defects in collagen biosynthesis and structure, human EDS has been reclassified into 13 subtypes based on major and minor criteria. Diagnosis of cEDS in humans relies on fulfilling minimal criteria (Table 2) as follows: Major criterion (1) skin hyperextensibility and atrophic scarring; plus either major criterion (2) generalized joint hypermobility and/or at least three minor criteria. Confirmatory molecular testing is obligatory to reach a final diagnosis.

In case 1, the affected dog met both major criteria. One minor criterion, skin fragility, was also noted in the case history. This animal was 9 months of age at the time of euthanasia – it is possible that had the animal survived into adulthood more of the minor criteria would have been met. Regardless, the diagnosis of cEDS in this case was confirmed by the demonstration of the missense mutation of the *COL5A1* gene.

In cases 2 and 3 the major criterion of skin hyperextensibilty (case 2 and 3) and atrophic scarring (case 2) was met, however joint hyper mobility was not noted. However, three minor criterion (easy bruising, traumatic splitting of the skin and family history of a first degree relative who meets clinical criteria) were fulfilled. The diagnosis of cEDS in these cases was confirmed by the demonstration of the missense mutation of the *COL5A1* gene.

Some confusion has existed as to terminology as ED-like syndrome in dogs has been previously termed cutaneous asthenia. The term cutaneous asthenia in humans is reserved for cases where abnormalities in both collagen and elastin (cutis laxa) exist. The cutis laxa syndromes are no longer included within the EDS classification system and have not been definitely diagnosed in animals.[18] Histopathology is typically pursued when heritable connective tissue disorders are suspected in animals. Morphologic changes in the skin may be absent or subtle on light microscopy, with reported changes typically including abnormal collagen fibers (collagen dysplasia) that are widely separated, finer and/or paler than normal, and haphazardly arranged. An increase in dermal elastic fibers has been described as a rare histopathologic feature of hereditary collagen dysplasia [41], but a distinction between the various forms of hereditary connective tissue disorders cannot be made based on morphologic assessment of the tissue. In most cases, ultrastructural examination and is required to confirm the collagen abnormality detected by light microscopy [41].

Histopathology of skin from the 3 affected dogs revealed dermal collagen dysplasia, with collagen fibers that varied in diameter and length and were haphazardly arranged. Case 1 also had an increase in dermal elastic fibers, multifocally; a VVG stain was not performed in cases 2 and 3. Cases 2 and 3 had more dermal fibroblasts and more space between collagen fibers, but this was not a feature of case 1. Dermal thickness of case 1 could not be compared to the control since the sections came from areas of the body with innate differences in dermal thickness, i.e. the back and abdomen, respectively. Cases 2 and 3 had normal dermal thicknesses.

Collagen protein analysis of case 1 showed an apparently normal collagen composition of the EDS skin compared to the control skin, with predominantly collagen I and smaller amounts of collagen III. Both EDS and control samples had, as expected, small amounts of collagen V, mostly in the pepsin-extracted crosslinked collagen fraction. There was no evidence of differentially solubility of any of the collagen fractions suggesting that the incorporation of all collagens into the dermal extracellular matrix was not impeded by the collagen V mutation. While not able to be quantified accurately, the EDS dermal pepsin extract appeared to contain a reduced amount of collagen V α1(V) chain relative to collagen I α1(I). This is consistent with the heterozygous *COL5A1* p.(Gly1013ValfsTer260) mutation characterized in case 1 which, because of the introduction of a premature termination codon would be expected to result in α1(V) mRNA instability (nonsense-mediated mRNA decay)[42]. This would lead to α1(V) haploinsufficiency. Since collagen V can only form as a protein heterotrimer of α1(V)_2_α2(V), this would result in an overall reduction in collagen V protein. A limitation of our studies is that we were unable to extract mRNA from the frozen tissue samples available, or obtain cell lines from case 1 to directly demonstrate the nonsense-mediated mRNA decay of the mutant *COL5A1* allele. Haploinsufficiency of *COL5A1* has been demonstrated in mice and humans and since the ratio of collagen V:collagen I is important for correct collagen I fibrillogenesis, reduced collagen V is associated with the EM findings of disorganized collagen fibrils and the clinical phenotype of cEDS [19, 21, 43].

Genetic analysis of cases 2 and 3 revealed a missense variant in the *COL5A1* gene, p.(Gly1571Arg). The predicted amino acid exchange occurs in the protein helical domain and by analogy with helix glycine substitution mutations in collagen I in osteogenesis imperfecta, collagen II in chondrodysplasias and collagen III and V in Ehlers Danlos syndrome, would be expected to disrupt collagen helix formation and/or stability.[44] In addition, intracellular retention of this structurally abnormal collagen could result in endoplasmic reticulum stress which can impact cellular function. In addition, the intracellular degradation of the mutant collagen by the cellular quality control machinery may lead to reduced secretion of collagen V containing the mutant α1(V), also resulting in reduced extracellular collagen V available to regulate collagen I fibrillogenesis. Furthermore, any secretion of the mutant collagen V may interfere with collagen interactions in the extracellular matrix.[44] Unfortunately neither dermal tissues, or cells, were available from these dogs to directly explore these pathological mechanisms.

Interestingly, in case 1 the variant was passed on from a clinically non-affected mosaic parent to the offspring. It is however unknown whether the mother was clinically completely normal or showed subtle signs of Ehlers-Danlos syndrome. The genetic analysis of case 1 and its parents confirmed the suspected de novo mutation event. Given that only one protein-changing *de novo* mutation event per generation is expected on average, this further strongly supports the causality of the variant for the EDS phenotype. We speculate that a similar independent *de novo* mutation event has happened in one of the parents of cases 2 and 3. Unfortunately, no DNA samples from their parents were available and we could not verify this assumption.

Our results are also consistent with recent findings in a cat that was also affected by Ehlers Danlos syndrome due to a de novo mutation in the COL5A1 gene.[45] Together these studies show that whole genome sequencing as part of a “precision medicine” approach can be a valuable tool to obtain a precise definition of the molecular lesion in domestic animal patients suspected to suffer from an inherited disease.

## CONCLUSION

To the best of our knowledge, we provide the first report of genetic variants in the *COL5A1* gene that are associated with the clinical presentation of EDS in dogs. Molecular screening by means of whole genome or whole exome sequencing or the targeted resequencing of a gene panel that includes the known EDS candidate genes from other species is a viable tool in the diagnosis of EDS in dogs. As in humans, any molecular techniques should always be combined with the established clinical criteria to confirm a diagnosis of EDS.

## ABBREVIATIONS

EDS: Ehler’s Danlos Syndrome
H&E: hematoxylin and eosin
MT: aMasson’s Trichome
VVG: Verhoeff van Gieson
BWA: Burrow-Wheeler Aligner
GATK: Genome Analysis Tool Kit
SNVs: single mucleotide variants
MQ: mapping quality
gVCF: genome variant call format
QD: quality of depth

## DECLARATIONS

### Ethics approval and consent to participate

All procedures for case 1 (FB) were approved by the Murdoch Children’s Research Institute Animal Ethics Committee (Approval # A815). Blood and skin samples for cases 2 and 3 (RA) were collected as part of routine veterinary diagnostic assessment.

### Consent for publication

Not applicable

### Availability of data and materials

- The datasets generated and/or analyzed during the current study are available in the European Nucleotide Archive (ENA), https://www.ebi.ac.uk/ena/data/search?query=SAMEA104091568+OR+SAMEA4867923

### Authors’ contributions

AB performed the genetic analysis and contributed to writing.

JB was involved in biochemical analysis of tissue samples, study design and contributed to writing the manuscript

SL was involved in biochemical analysis of tissue samples and contributed to writing the manuscript

EH conducted the transmission electron microscopy

SK was involved in interpretation of histology for case 1 and contributed to writing.

MY was involved in case management, tissue collection and clinical images

AR identified the cases from Morocco and was involved in clinical management and sample acquisition.

VJ performed the bioinformatics processing of the whole genome sequence data and handled sequence data deposition to the ENA.

MW performed the histopathological analyses on the cases from Morocco and contributed to writing the manuscript.

TL supervised the genetic analysis and contributed to manuscript writing.

FB was involved in clinical case management, sample collection, major contributor in writing manuscript

## Acknowledgements

The Next Generation Sequencing Platform of the University of Bern is acknowledged for performing the whole genome re-sequencing experiment and the Interfaculty Bioinformatics Unit of the University of Bern for providing computational infrastructure. We acknowledge the Dog Biomedical Variant Database Consortium (Gus Aguirre, Catherine André, Danika Bannasch, Doreen Becker, Brian Davis, Cord Dröegemüller, Kari Ekenstedt, Kiterie Faller, Oliver Forman, Steve Friedenberg, Eva Furrow, Urs Giger, Christophe Hitte, Marjo Hytönen, Vidhya Jagannathan, Tosso Leeb, Hannes Lohi, Cathryn Mellersh, Jim Mickelson, Anita Oberbauer, Jeffrey Schoenebeck, Kim Summers, Frank van Steenbeck, Claire Wade) for sharing whole genome sequencing data from control dogs and wolves. We thank all owners for donating samples, and additional data of their animals.

